# Differential recognition of computationally optimized H3 hemagglutinin influenza vaccine candidates by human antibodies

**DOI:** 10.1101/2022.02.24.481830

**Authors:** Nada Abbadi, Kaito Nagashima, Alma Pena-Briseno, Ted M. Ross, Jarrod J. Mousa

**Affiliations:** Department of Infectious Diseases, College of Veterinary Medicine, University of Georgia, Athens, GA, USA; Center for Vaccines and Immunology, College of Veterinary Medicine, University of Georgia, Athens, GA, USA; Department of Biochemistry and Molecular Biology, Franklin College of Arts and Sciences, University of Georgia, Athens, GA, USA

## Abstract

Among circulating influenza viruses in humans, H3N2 viruses typically evolve faster than other subtypes and have caused severe illness and deaths in millions of people since emerging in 1968. Computationally optimized broadly reactive antigen (COBRA) technology is one strategy to broaden vaccine-elicited antibody responses among influenza subtypes. In this study, we determined the structural integrity of an H3N2 COBRA HA, TJ5, and, as nearly all humans have pre-existing immunity to H3N2 influenza viruses, we probed the antigenic profile of several H3N2 COBRA HAs by assessing recognition of these immunogens by human B cells and monoclonal antibodies (mAbs). Of three recently described COBRA H3 HA antigens (TJ5, NG2, and J4), we determined that TJ5 and J4 HA proteins recognize pre-existing B cells (from the 2017-2018 vaccine season) more effectively than NG2 HA and a wild type Hong Kong/4801/2014 protein. H3 HA-specific human mAbs recognize wild type and COBRA HA proteins, and have functional activity against a broad panel of H3N2 viruses. mAb TJ5-5 recognizes TJ5 and J4 HA proteins, but has poor recognition of NG2 HA, similar to the global B cell analysis. To probe these recognition differences and to verify the structural integrity of the TJ5 HA protein, we determined a 3.4 Å structure via cryo-electron microscopy of TJ5-5 complexed with the TJ5 HA, which revealed residues important to the differential binding. Overall, these studies determined that COBRA H3 HA proteins have correct antigenic and structural features, and are recognized by B cells and mAbs isolated from seasonally vaccinated humans.

**Importance:** Vaccine development for circulating influenza viruses, particularly for the H3N2 subtype, remains challenging due to consistent antigenic drift. Computationally optimized broadly reactive antigen (COBRA) technology has proven effective for broadening influenza hemagglutinin (HA) elicited antibody responses compared to wild type immunogens. Here we determined the structural features and antigenic profiles of H3 COBRA HA proteins. Two H3 COBRA HA proteins, TJ5 and J4, are better recognized by pre-existing B cells and monoclonal antibodies from the 2017-2018 vaccine season compared to COBRA NG2 and a wild type A/Hong Kong/2014 HA protein. We determined a cryo-EM structure of one mAb that poorly recognizes NG2, mAb TJ5-5, in complex with the TJ5 COBRA HA protein and identified residues critical to mAb recognition. As NG2 is more effective than TJ5 for a recent Hong Kong/2019 virus, these data provide insights into the diminished effectiveness of influenza vaccines across vaccine seasons.

## Introduction

Influenza virus is a global respiratory pathogen that has been circulating in humans for hundreds of years^1^. The World Health Organization (WHO) estimates annual influenza epidemics cause 2–5 million severe cases and 250,000 to 500,000 deaths^2^. Influenza disease symptoms range from mild fever, sore throat, coughing, nasal discharge, headache, and myalgia to more severe cases that can lead to the development of bronchitis or pneumonia^3^. There are four types of influenza viruses, including A, B, C and D, of which types A and B cause seasonal epidemics in humans^4^. The current circulating influenza A subtypes are H1N1 and H3N2, and these subtypes have caused three of the four influenza pandemics in the last century, including the 1918 H1N1 “Spanish flu” pandemic^1^, the 1968 H3N2 “Hong Kong flu” pandemic^5^, and the 2009 H1N1 “Swine flu” pandemic^6^. H3N2 influenza viruses evolve more rapidly than H1N1 viruses, which results in more frequent updates to the H3N2 strain used in seasonal influenza vaccines^7^, and the WHO has recommended 28 vaccine strain changes since their introduction^8^.

On the surface of influenza virus, the hemagglutinin (HA) and neuraminidase (NA) proteins are the major targets of protective antibodies. HA is responsible for binding to sialic acid found on the host cell surface, allowing viral entry to the cell^9^. NA cleaves the sialic acid residues at the end of the influenza life cycle in the cell, allowing the virus to be released from the infected cell resulting in viral spread to neighboring uninfected cells^10^. Aside from its enzymatic action in releasing the virus, NA can also bind to sialic acid and assist in viral entry^11^. HA is the most abundant protein on the surface of the virion and is immunodominant, which makes it the target of all current licensed influenza vaccines^12,13^. However, virus evolution and immune pressure cause frequent point mutations resulting in changes to the HA antigenic structure, facilitating evasion of the immune response.

The most effective countermeasure against influenza disease is vaccination. There are three classes of licensed seasonal vaccines: inactivated, live attenuated, and recombinant HA vaccines^14^. All of these vaccines contain a mixture of viruses representing both influenza A (H1N1 and H3N2 subtypes) and B (Victoria and Yamagata lineages) viruses^15^. Due to the mutable nature of influenza viruses, antibodies generated from a previous infection or vaccination may fail to recognize the drifted virus, leading to influenza epidemics^16^. This constant change in the antigenicity of influenza makes it difficult to maintain protective immunity against the virus. Therefore, there is a need for a universal influenza vaccine that protects against multiple strains of a given subtype or even across subtypes to decrease the possibility of another deadly pandemic. Computationally optimized broadly reactive antigen (COBRA) proteins are a tool to broaden vaccine-elicited antibody reactivity by incorporating influenza HA (or NA) sequences from multiple influenza seasons to generate a consensus antigen. COBRA antigens have been successful in expanding the breadth of vaccine-elicited antibody responses for H1N1^17,18^, H3N2^19,20^, H2N2^21^, and H5N1^22,23^ influenza viruses, and have recently been applied to NA^24^.

Next-generation H3 HA COBRA antigens have recently been developed, and differing immune responses compared to previous H3 COBRA HAs was observed^25^. For example, while COBRA TJ5 HA elicited increased antibody breadth, the effectiveness was limited against a more recent A/Hong Kong/45/2019 (HK19) virus compared to a next-generation H3 COBRA NG2 (*in press, JVI)*. To determine the differential mechanism of COBRA mediated breadth, we probed B cell and human monoclonal antibody (mAb) recognition of COBRA HAs from subjects vaccinated with the 2017-2018 split-inactivated seasonal influenza vaccine. Pre-existing B cells from these subjects have higher recognition of COBRA TJ5 and J4 HAs compared to NG2 and Hong Kong/4801/2014 HA proteins. All mAbs bind to the receptor binding site, except for one mAb targeting the stalk domain, which verified the antigenic integrity of the recombinant COBRA HA immunogens. We discovered that one mAb, TJ5-5, has diminished binding to COBRA HA NG2 compared to COBRA HA TJ5 similar to the B cell screening. Structural information of COBRA HA immunogens is limited, with the exception of the structure of an H5 COBRA HA protein^26^. To determine the mechanism of diminished recognition and to verify the structural integrity of the H3 COBRA antigen, we determined a cryo-EM reconstruction of mAb TJ5-5 in complex with COBRA TJ5 HA and we present the structural basis for mAb recognition.

## Materials and Methods

### Human subject samples

All human studies were approved by the University of Georgia Institutional Review Board. mAb isolation was conducted from subjects vaccinated with the 2017-2018 seasonal influenza vaccine (Fluzone) from peripheral blood mononuclear cells (PBMCs) isolated from blood draws 21-28 days following vaccination.

### B cell expansion of human subject PBMCs

PBMCs were plated at a density of 25,000 cells/well in a 96-well plate on a layer of gamma-irradiated NIH 3T3 cells (20,000 cells/well) expressing hCD40L, hIL-21, and hBAFF in the presence of CpG and cyclosporine A as previously described^27,28^. B cell supernatants were screened by enzyme-linked immunosorbent assay (ELISA) at 7 days post-plating of PBMCs.

### Expression and purification of recombinant influenza HA proteins

Trimeric wild-type HA or COBRA HA ectodomains were expressed and purified in Expi293F cells following the manufacturer’s protocol and as previously described^29^. Collected supernatants containing the HA antigens were purified on a HisTrap Excel column following the manufacturer’s recommended protocol. Eluted fractions were pooled and purified proteins were verified for integrity by probing with an anti-HIS tag antibody (Biolegend) as well as with subtype-specific mAbs via SDS-PAGE and Western blot.

### ELISA screening of B cells, hybridoma supernatants, and mAbs

384-well plates (VWR) were coated with recombinant HA proteins diluted to 2 μg/mL in PBS at 4 °C overnight. Plates were washed once with water, then blocked with 2% blocking buffer (PBS + 2% non-fat dry milk (Bio-Rad) + 2% goat serum + 0.05% Tween-20) for 1 hr at room temperature. Plates were washed three times with water, and 25 μL of B cell supernatants, hybridoma supernatants, or mAbs were added. mAbs were serially diluted three-fold in PBS from 20 μg/mL prior to addition for twelve total dilutions. Plates were incubated at 37 °C for 1 hr, then washed three times with water. Goat anti-human IgG Fc-AP secondary antibody (Southern Biotech), diluted 1:4000 in 1% blocking buffer (1:1 dilution of PBS and 2% blocking buffer), was added and plates were incubated at room temperature for 1 hr. Plates were then washed five times with PBS-T (PBS + 0.05% Tween-20). *p*-Nitrophenyl phosphate (PNPP) substrate, diluted in substrate buffer (1.0 M Tris + 0.5 mM MgCl_2_, pH=9.8) to 1 mg/mL, was added, and plates were incubated for 1 hr and read at 405 nm on a BioTek plate reader. The EC_50_ value for each mAb was determined by using the four-parameter logistic curve fitting function in GraphPad Prism software.

### Generation of HA-reactive mAbs

Eight days following plating of PBMCs, wells identified to contain positive B cells by ELISA were selected for electrofusion to generate hybridomas as previously described^27,28^. Hybridomas were plated in 384-well plates for HAT selection, and grown for 14 days at 37°C, 5% CO_2_. Following screening by ELISA, hybridomas were single-cell sorted using a MoFlo Astrios cell sorter using live/dead staining by propidium iodide. The sorted hybridomas were cultured in 25% Media E (StemCell) + 75% Media A (StemCell) for two weeks, then subjected to another round of screening by ELISA. Hybridomas with the highest signal were grown in 250 mL serum-free media (Gibco) for approximately one month. Secreted mAbs were purified using a Protein G column (GE Healthcare) and concentrated for use in downstream assays.

### Hybridoma sequencing

Hybridoma cell lines encoding each mAb were sequenced utilizing the primers described by Guthmiller *et al.*^29^. Briefly, RNA was extracted from each hybridoma and cDNA was generated using the SuperScript IV First-Strand cDNA Synthesis Kit (Invitrogen). A nested PCR protocol was used to generate sequencing products. In the first nested PCR step, a primer mix specific to the heavy, kappa, or lambda chain *V* gene and the constant region were used to amplify the variable region using the cDNA as template. In the second PCR step, the first PCR product was used as a template with a nested primer mix to improve product specificity and yield. The second nested PCR products were sequenced using the constant region primer and the *V, D,* and *J* alleles were identified by IMGT/V-QUEST^30^.

### Hemagglutination inhibition assay

The HAI titer for each mAb was determined as previously described^31^. Influenza viruses were titered to eight HAUs (hemagglutination units). mAbs diluted to 50 μg/mL in PBS were added to the first well of a 96-well U-bottom plate (VWR), and diluted five-fold in PBS. Eight HAUs of virus with 40 nM Osteltamivir were added in a 1:1 ratio to each mAb dilution, and each well was mixed and incubated for 30 min at room temperature. Following this, 50 μL of 0.8% Guinea pig red blood cells (Lampire) were added per well. Plates were read 1 hour after the addition of 0.8% Guinea pig red blood cells.

### Focal reduction assay

Focal reduction assays (FRAs) were completed for each mAb as previously described^31^. MDCK cells were plated in 96-well plates overnight to achieve >95% confluency the next day. Cells were washed twice with PBS, and 50 μL of virus growth media (VGM: DMEM + 2 μg/mL TPCK-trypsin + 7.5% BSA) were added and the plates were returned to the incubator at 37°C, 5% CO_2_. mAbs at 50 μg/mL were serially diluted five-fold in VGM, and virus was diluted to a concentration of 1×10^4^ FFU/mL in VGM. MDCK cells were washed with PBS and 25 μL serially diluted mAbs were added, followed by 25 μL of 1×10^4^ FFU/mL of virus. Plates were incubated at 37 °C, 5% CO_2_ for 2 hr, and then 100 μL/well of overlay media (1.2% Avicel + modified Eagle media (MEM) + 7.5% BSA) were added and incubated overnight. The overlay was removed and wells were washed twice with PBS. Ice-cold fixative (20% formaldehyde + 80% methanol) was added and plates were incubated at 4 °C for 30 min. Plates were washed twice with PBS and permeabilization buffer (PBS + 0.15% glycine + 0.5% Triton-X 100) was added, followed by a 30 min incubation. Plates were washed three times with PBS-T and primary IAV anti-NP mouse antibody (IRR), diluted 1:2000 in ELISA buffer (PBS + 10% goat serum + 0.1% Tween-20), was added. Plates were incubated at room temperature for 1 hr. Plates were then washed three times with PBS-T and secondary goat anti-mouse IgG-HRP antibody (Southern Biotech), diluted 1:4000 in ELISA buffer, was added. Plates were incubated at room temperature for 1 hr and then washed with PBS-T. KPL TrueBlue Peroxidase substrate was added per well and plates were incubated for 10-20 min. Plates were washed, dried, and foci were enumerated using an ImmunoSpot S6 ULTIMATE reader with ImmunoSpot 7.0.28.5 software (Cellular Technology Limited). Neutralizing IC50s were calculated using the GraphPad Prism four-parameter logistic curve fitting function.

### Epitope binning by biolayer interferometry

The panel of mAbs isolated from human subjects were competed for binding using the A/Hong Kong/4801/2014 HA protein on the OctetRED384 system as previously described^32^. Anti-penta-HIS biosensors (Sartorius) were immersed in kinetics buffer (PBS + 0.5% BSA + 0.05% Tween-20) for 60 s to obtain a baseline reading. Biosensors were then loaded with 100 μg/mL of A/Hong Kong/4801/2014 HA protein diluted in kinetics buffer for 60 secs. Biosensors were returned to kinetics buffer for a baseline of 60 s. Following this, biosensors were immersed in the first mAb (100 μg/mL in kinetics buffer) for 300 s for the association step. The biosensors were then immersed in the competing, second mAb (100 μg/mL in kinetics buffer) for 300 s. The biosensors were then regenerated in 0.1 M glycine, pH = 2.7 and PBS alternately for three cycles before proceeding to the next mAb competition set. The extent of competition was calculated as the percentage of the signal from the second mAb in the second association step in the presence of the first mAb to that of the second mAb alone in the first association step for all biosensors. A ratio of <=31 was considered complete competition, >31 and <=70 moderate competition, and >70 no competition.

### Electron microscopy of the TJ5+TJ5-5 complex

TJ5 HA and Fab were mixed in a 1:2 ratio, and complexes were purified by size exclusion chromatography on a Superdex S200, 16/600 column (GE Healthcare Life Sciences) in 20 mM Tris pH 7.5, 100 mM NaCl. Quantifoil 1.2/1.3 400 mesh copper grids were glow discharged for 45 seconds at 25 mAmp current on the carbon side. Grids for cryo-EM were prepared by applying protein (1.2 mg/mL) to grids with a blotting time of 8 or 10 seconds, at 4 degrees C with 100% humidity, force=3, and 10s wait time. Grids with appropriate ice thickness were used for data collection on a Thermo Fisher Glacios 200 kV microscope with a Falcon 4 direct electron detector at 190,000x magnification, an image pixel size of 0.526 Å, a dose of 57.22 e-/ Å^2^, and 30 frames per movie. Data were analyzed in cryoSPARC for patch motion correction, patch CTF correction, particle picking and extraction, multiple rounds of 2D and 3D class averaging, and refinement. The structure was manually built in COOT.

## Results

### Oligoclonal B cell recognition of COBRA HA proteins

To determine the antigenic features of H3 COBRA HA proteins and the ability to interact with B cells from previous influenza exposure, we stimulated B cells derived from subjects vaccinated with the split inactivated 2017-2018 seasonal influenza vaccine (Fluzone), which contained A/Hong Kong/4801/2014 (HK14) as the H3 vaccine component. B cells were stimulated for six days on an irradiated feeder layer expressing human CD40L, human IL-21, and human BAFF in the presence of CpG. Stimulated B cell supernatant reactivity was assessed by ELISA after six days for eight human subjects. The antibody response from stimulated B cell supernatants was measured using H3 HA protein from the seasonal strain HK14 and H3 COBRA HAs TJ5, J4 and NG2. TJ5 was developed from HA sequences spanning 2008 – 2012, J4 from 2013 – 2016, and NG2 from 2016 – 2018. In the majority of subjects, reactivity to the TJ5 and J4 HA proteins generally had the highest response, and these signals were consistently higher than the NG2 HA protein and HK14 HA proteins, suggesting that TJ5 and J4 H3 COBRA HA proteins are able to better recall previous influenza virus exposures (prior to 2017-2018) than NG2 and HK14 HAs **(Figure 1)**. Among the COBRA HA proteins, TJ5 HA shares higher sequence identity with J4 compared to NG2 and HK14. This along with the different time frame used to generate each COBRA may explain the differences in antibody titers detected against these antigens.

**Figure 1.**
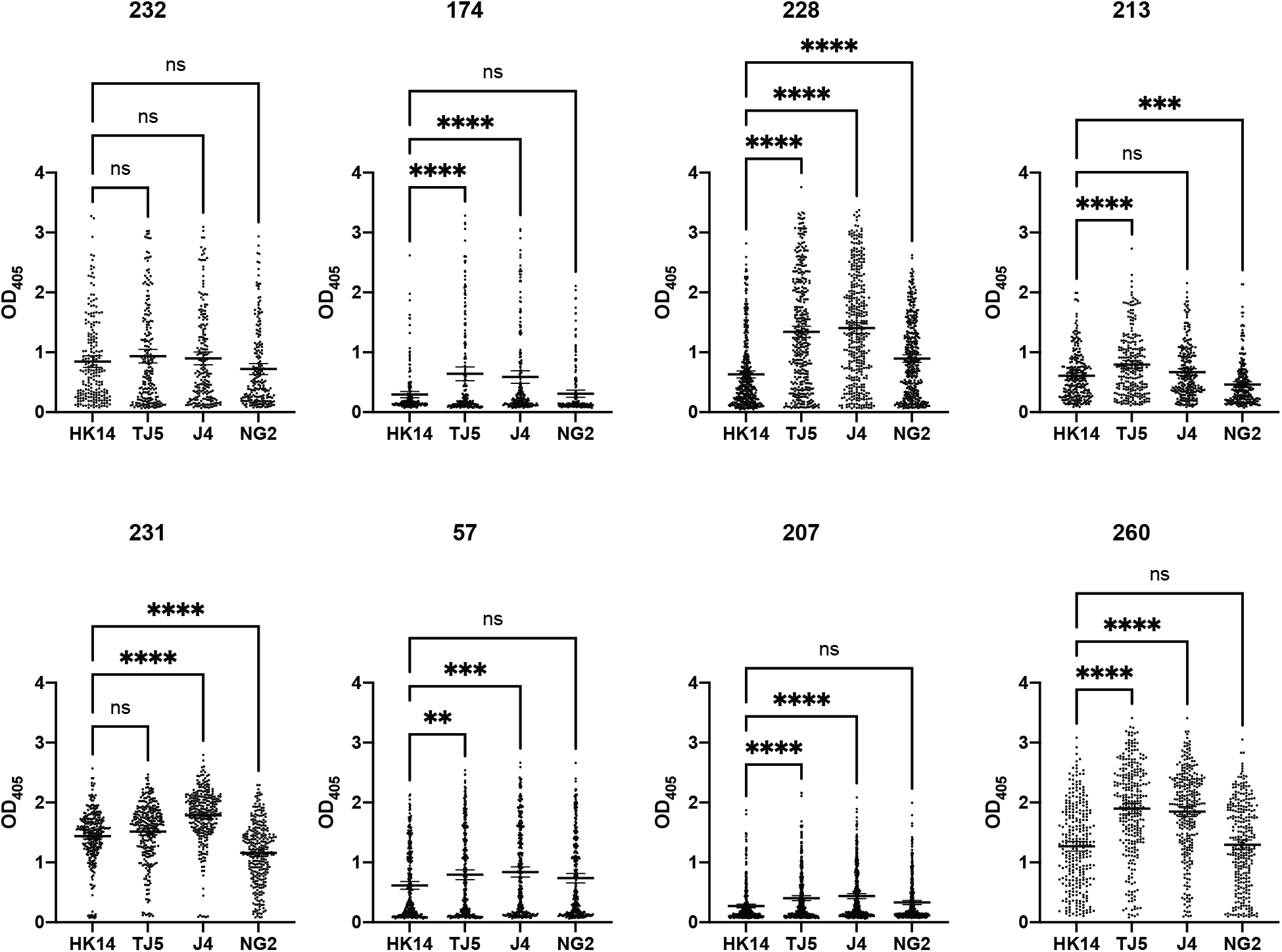
Binding titers of oligoclonal B cell supernatants 21 days post-vaccination from eight human subjects. The optical density determined by ELISA is shown for HK14 HA, TJ5 COBRA HA, J4 COBRA HA, and NG2 COBRA HA proteins for B cells stimulated from subjects receiving the 2017-2018 QIV. Supernatants from stimulated PBMCs were screened by ELISA using plates coated with the indicated antigen. PBMCs were standardized to 25,000 cells per well. Each circle indicates 1 well. ****P<0.0001, ***P=0.0002-0.0006, **P=0.0047, ns=not significant (P>0.05).

### Human mAb isolation and binding characterization

To further probe the reactivity of pre-existing human B cells and the antigenicity of the COBRA HA proteins, we isolated twelve human mAbs using the TJ5 COBRA HA as a baiting antigen since it was among the proteins that had the highest reactivity to pre-existing B cells. mAbs were sequenced, and 75% of mAbs utilize the commonly used VH1-69 heavy chain V gene, while the remaining mAbs utilized VH1-2, VH3-21, and VH4-39 genes **(Figure S1A, Table S1, S2).** All VH1-69 utilizing mAbs were paired with VL1-40 light chain. The lengths of the heavy and light chain junctions ranged from 11-19 amino acids for the heavy chain and 10-19 amino acids for the light chain **(Figure S1B)**. The percent identities of the V genes to the germline sequence had an average of 89% for the heavy chain and 96% for the light chain **(Figure S1C)**.

We next determined the reactivity of the mAbs by ELISA to the COBRA HA proteins as well as to several HA proteins derived from wild type influenza viruses (**Figure 2)**. All mAbs had some reactivity to all antigens tested, while mAbs TJ5-2, TJ5-4, TJ5-5, TJ5-6, TJ5-8, TJ5-9, TJ5-13, and TJ5-15 had the lowest EC_50_ values (indicating stronger binding) across all HA proteins. mAb TJ5-13 had consistently high binding affinity across all HA proteins tested, while mAbs TJ5-1, TJ5-11, and TJ5-14 had the lowest affinity. When assessing binding to each HA protein, binding of mAbs was overall better for TJ5 HA, J4 HA, and pre-2014 wild type HA proteins compared to NG2 HA and HK14 HA proteins, similar to our data observed in the oligoclonal B cell screens **(Figure 2).**

**Figure 2.**
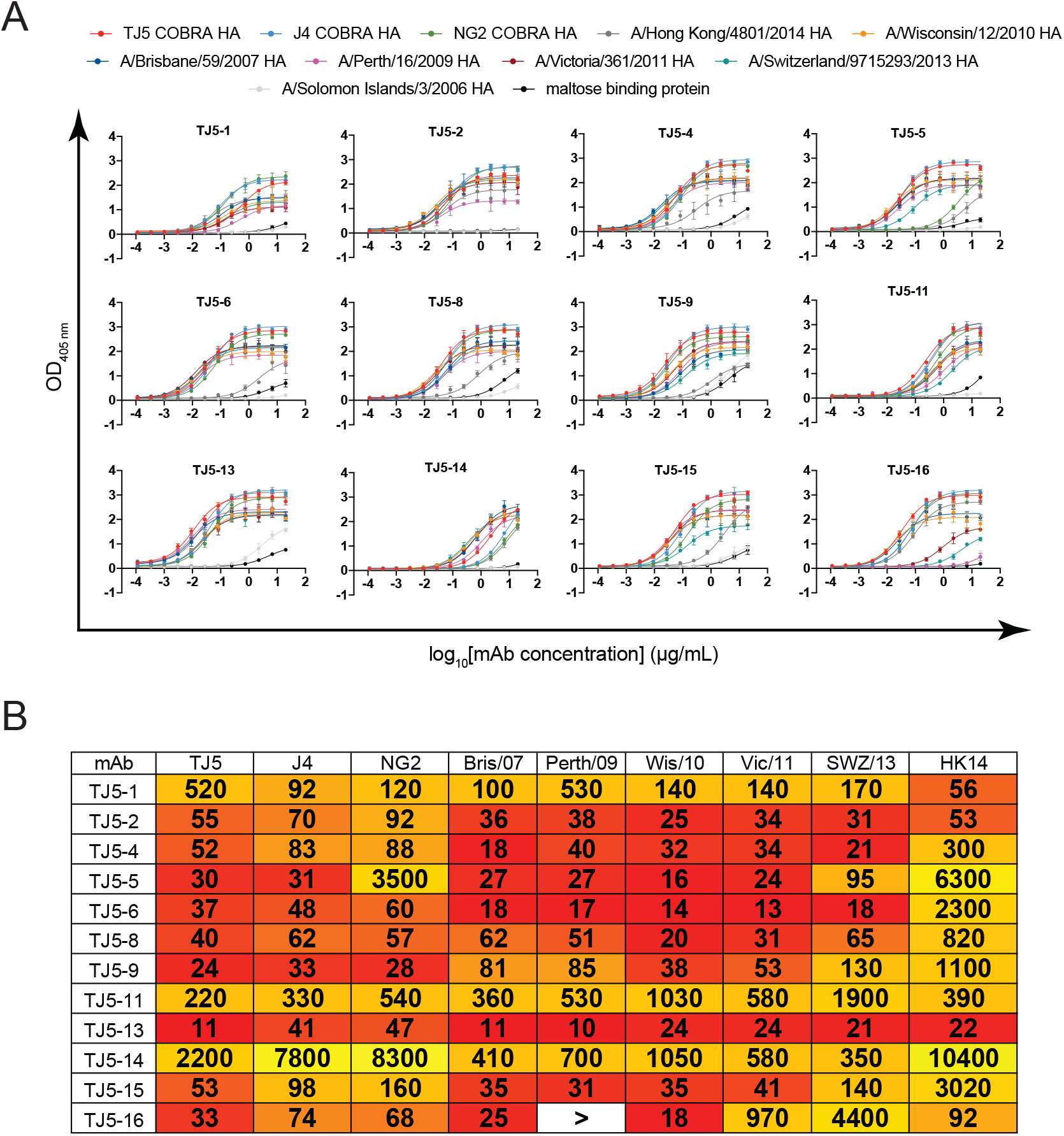
Reactivity of mAbs isolated from 2017-2018 QIV-vaccinated subjects. (A) ELISA binding curves for each mAb across a panel of antigens. Each data point is the average of four replicates and error bars indicate the standard deviation. ELISAs were completed with each mAb serially diluted three-fold. Data points are colored according to the legend at the top of the graphs. (B) Half-maximal effective concentrations (EC_50s_) to each antigen are displayed for each mAb. The heat map is colored from red (low EC_50_, high binding) to yellow (high EC_50_, low binding). > indicates the signal at 20 μg/mL did not reach 1.5, or the calculated EC_50_ was outside the tested concentration range due to an overall low signal.

### Functional analysis of isolated H3 reactive mAbs

In addition to binding analysis, we characterized the functional activity of the 12 isolated mAbs using HAI and neutralization assays **(Figure 3).** In our experiments we used a panel of seasonal H3N2 influenza viruses from 1968 – 2017. All mAbs except TJ5-1 and TJ5-2 had HAI activity against at least three viruses, and several mAbs (TJ5-4, TJ5-6, TJ5-15) had HAI activity against H3N2 viruses ranging from 2004-2014 **(Figure 3A)**. To further measure the functionality of these mAbs, we tested the neutralization activity against a panel of H3N2 viruses through focal reduction assays **(Figure 3B, 3C).** The majority of mAbs were able to neutralize nearly all viruses **(Figure 3C)**, with TJ5-13 having potent neutralizing activity against all viruses tested from 2004 to 2017. Despite its lack of HAI activity, TJ5-1 had the broadest neutralizing activity against the panel of viruses, which indicates that this mAb is functional against influenza virus using an alternative mechanism to hemagglutination inhibition. In contrast, TJ5-2 was non-neutralizing. These data demonstrate that the majority of the isolated mAbs that could potentially be recalled with COBRA HA vaccination were functional against a large panel of seasonal H3N2 influenza viruses.

**Figure 3.**
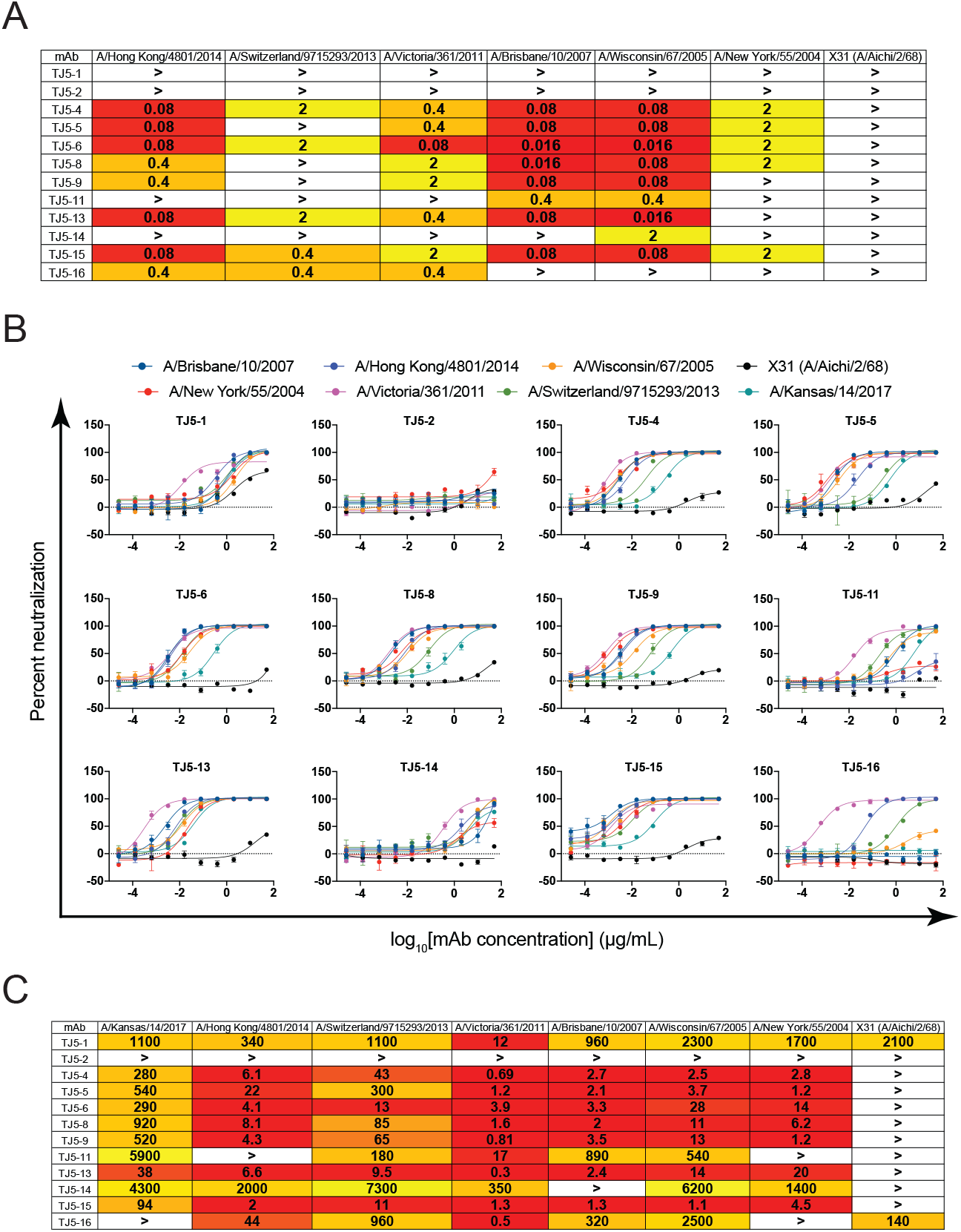
Functional properties of mAbs isolated from 2017-2018 QIV-vaccinated subjects. (A) HAI inhibitory titers for each mAb against a panel of H3N2 influenza viruses. > indicates no HAI activity was seen at 2 μg/mL or below. Data are the average of two replicates. (B) Focus reduction assay curves for each mAb against a panel of H3N2 viruses. Data are colored according to the legend. Each data point represents the average of three replicates, and error bars are the standard deviation. (C) Half-maximal inhibitory concentrations of each mAb are shown for a panel of H3N2 viruses determined from curves in (B). In (A) and (C), data are colored from high HAI or FRA activity (red) to low (yellow). > indicates mAbs did not cause 50% neutralization at the highest concentration tested, or the IC50 was outside the range of the curve due to low neutralization.

### Epitope binning analysis for the panel of H3 HA specific human mAbs

HA specific antibodies can either target the head domain or the stalk domains. The HA head is immunodominant upon natural infection or vaccination, while the stalk domain is subdominant^33^. To further characterize the isolated mAbs, we performed epitope binning experiments to determine the specific epitopes on the HA structure targeted by these mAbs **(Figure 4)**. We used biolayer interferometry to compete the isolated mAbs against previously published H3 HA-specific mAbs with known epitopes on the HA protein. Five control mAbs binding to different epitopes on HA were used in this experiment: mAbs C05 and F045-92 target the receptor binding site (RBS) on the HA^34,35^; mAb CR8020 binds to the HA stem^36^; mAb C585 binds to the receptor binding subdomain^37^; and mAb F005-126 binds to a distinct epitope interacting with site L (residues 171 to 173, 239, and 240) and site R (residues 91, 92, 270 to 273, 284, and 285) as well as the glycan linked to N285^38^ **(Figure 4A).** All isolated mAbs except TJ5-1 competed with the RBS mAbs C05 and F045-92, indicating that the RBS is the target epitope for these mAbs. TJ5-1 competed with the stem antibody CR8020 **(Figure 4B),** which correlated well with our data demonstrating that TJ5-1 had broad neutralizing activity but no HAI activity due to its targeting of the conserved stalk domain. Overall, these data indicate the COBRA H3 HA proteins retain antigenic properties of wild type viruses, and can potentially recall both head and stalk domain targeting B cells.

**Figure 4.**
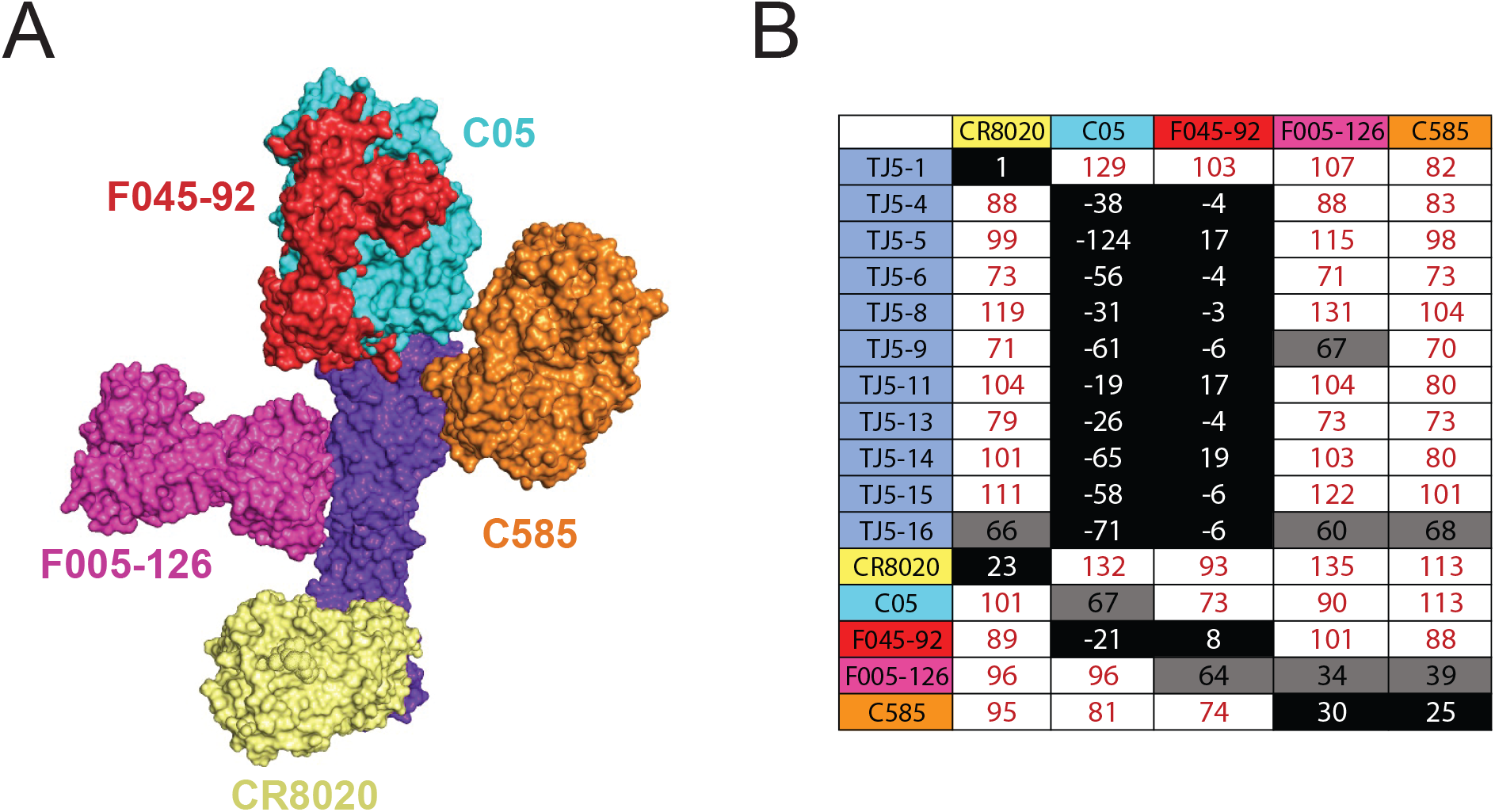
Epitope binning for the panel of H3 HA specific human mAbs. (A) Model of HK14 HA in complex with the five control antibodies used for epitope binning C05 (PDB 4FP8), F045-92 (PDB 4O58), C585 (6PDX), F005-126 (PDB 3WHE) and CR8020 (PDB 3SDY).(B) Epitope binning was performed with HK14 HA protein. Competition was measured as the percentage of the response from the association of the second antibody (horizontal axis) in the presence of the first antibody (vertical axis) as compared to the second antibody alone. Black indicates complete competition, gray moderate competition, and white no competition. Control mAbs are colored according to their position in (A).

### Structural basis for COBRA recognition

A recent study showed that vaccination with the H3 COBRA TJ5 did not elicit HAI+ antibodies against the antigenically drifted A/Hong Kong/45/2019 H3N2 virus. Among the isolated mAbs, TJ5-5 showed potent binding to TJ5 and J4 HA proteins, but poor binding to NG2 and HK14 HA proteins. HK19 is a future drift strain compared to the COBRA immunogens, yet has a very similar sequence to NG2 with approximately 98% identity. To determine the structural correlates of decreased NG2 recognition in our B cell screening and by mAb TJ5-5, we determined the structure of the TJ5-5 Fab in complex with TJ5 HA by cryo-electron microscopy to a resolution 3.4 Å **(Figure 5A, 5B, S2, Table S3).** The overall structure of the COBRA TJ5 HA aligns very well to wild type H3 HA structures of Hong Kong/68, Victoria/2011, and Switzerland/2013 HA proteins, with RMSDs of 0.748, 0.6, and 0.624, respectively **(Figure 5C)**. The amino acid residues on the HA receptor binding site Y98, N145, S137, K189, D190, N225 and R222 were found to have electrostatic interactions important to binding with the heavy chain residues Y36, F62, M59, N109, E82 and T29, respectively **(Figure 5D)**. In the light chain, one amino acid interaction was observed between N158 on HA and Y38 on the LC **(Figure 5E).** This shows that the heavy chain plays the more dominant role in HA recognition.

**Figure 5.**
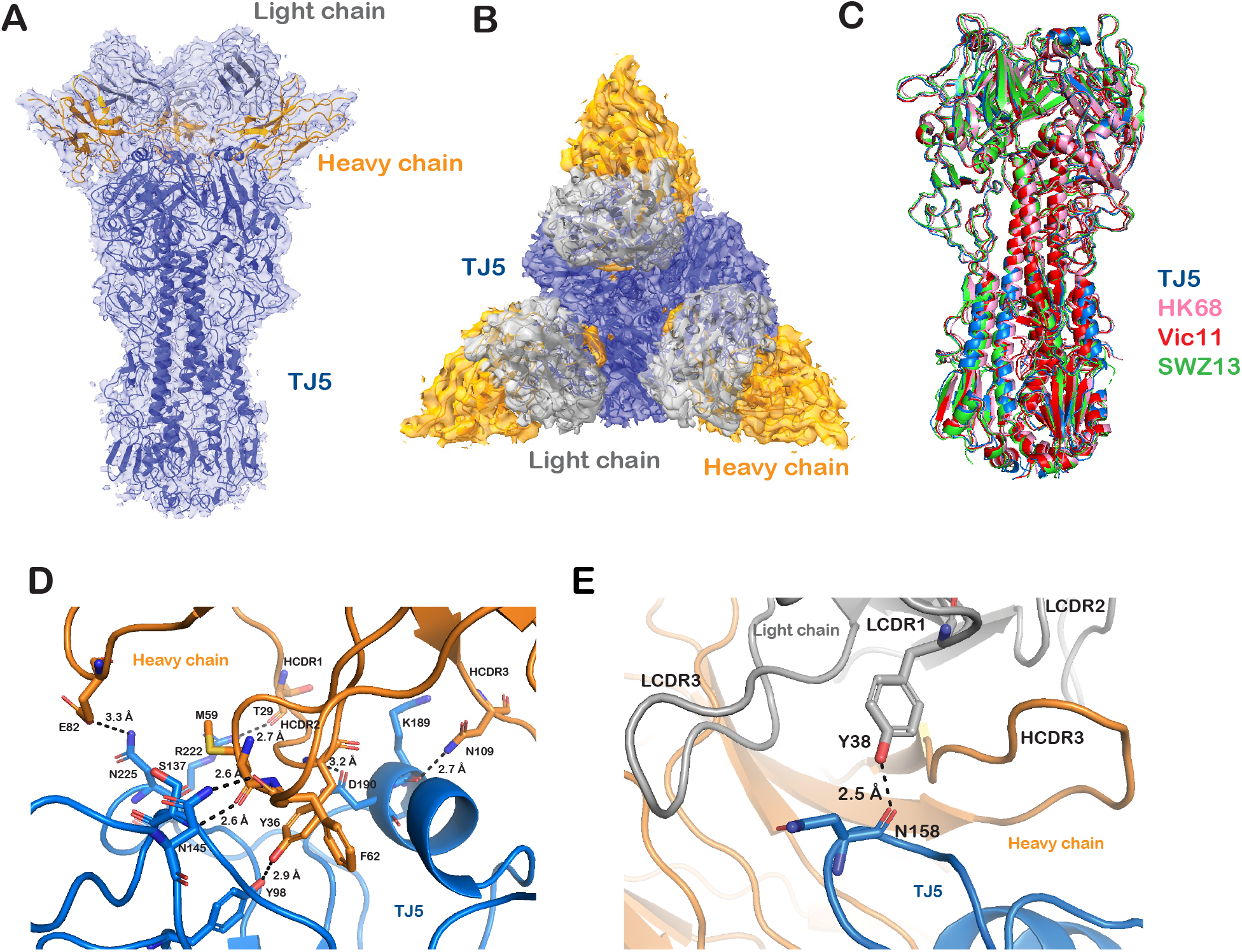
Cryo-EM structure of COBRA TJ5 complexed with mAb TJ5-5. A) Structure of TJ5 HA trimer bound to TJ5-5. B) Top view of TJ5-fab complex. C) Alignment of H3 HA molecule from TJ5 (Blue), A/Hong Kong/1/1968 (Pink) RMSD= 0.748, A/Victoria/361/2011 (Red) RMSD= 0.6 and A/Switzerland/9715293/2013 (Green) RMSD= 0.624. D) shows the amino acid interactions between the heavy chain and HA binding site. E) shows the amino acid interactions between the light chain and the HA binding site. The heavy chains are shown as orange, the light chains are shown in grey and the HA is shown in blue. The CDR loops for both heavy and light chains are displayed in the figures.

Multiple amino acid differences located near the binding site of mAb TJ5-5 were found in HA sequences of NG2, SWZ/13 and HK/19 when compared with the TJ5 sequence, including T128A, S137F, A138S, R142G, N144S, N145S, F159Y and N225D **(Figure 6A).** We determined that the lack of NG2 recognition and SWZ/13 neutralization could be due to the mutations N145S and N225D which can prevent binding to F62 and E82 residues on the HC, respectively **(Figure 6B, 6C).** In addition to N145S and N225D differences, S137F is present in HK/19 HA, which could also affect the lack of HAI activity observed for TJ5 vaccination against HK/19 (*in press, JVI)* **(Figure 6C).** Other mAbs such as TJ5-9, TJ5-13 and TJ5-15 showed potent NG2 binding as well as strong neutralization against SWZ/13, which could mean that these mAbs utilize different amino acid residues to bind to HA that are not mutated. This structural analysis indicates that antigen recognition and neutralization of influenza viruses by COBRA HA-targeting antibodies are largely dependent on the presence of a small number amino acids on the COBRA HA proteins.

**Figure 6.**
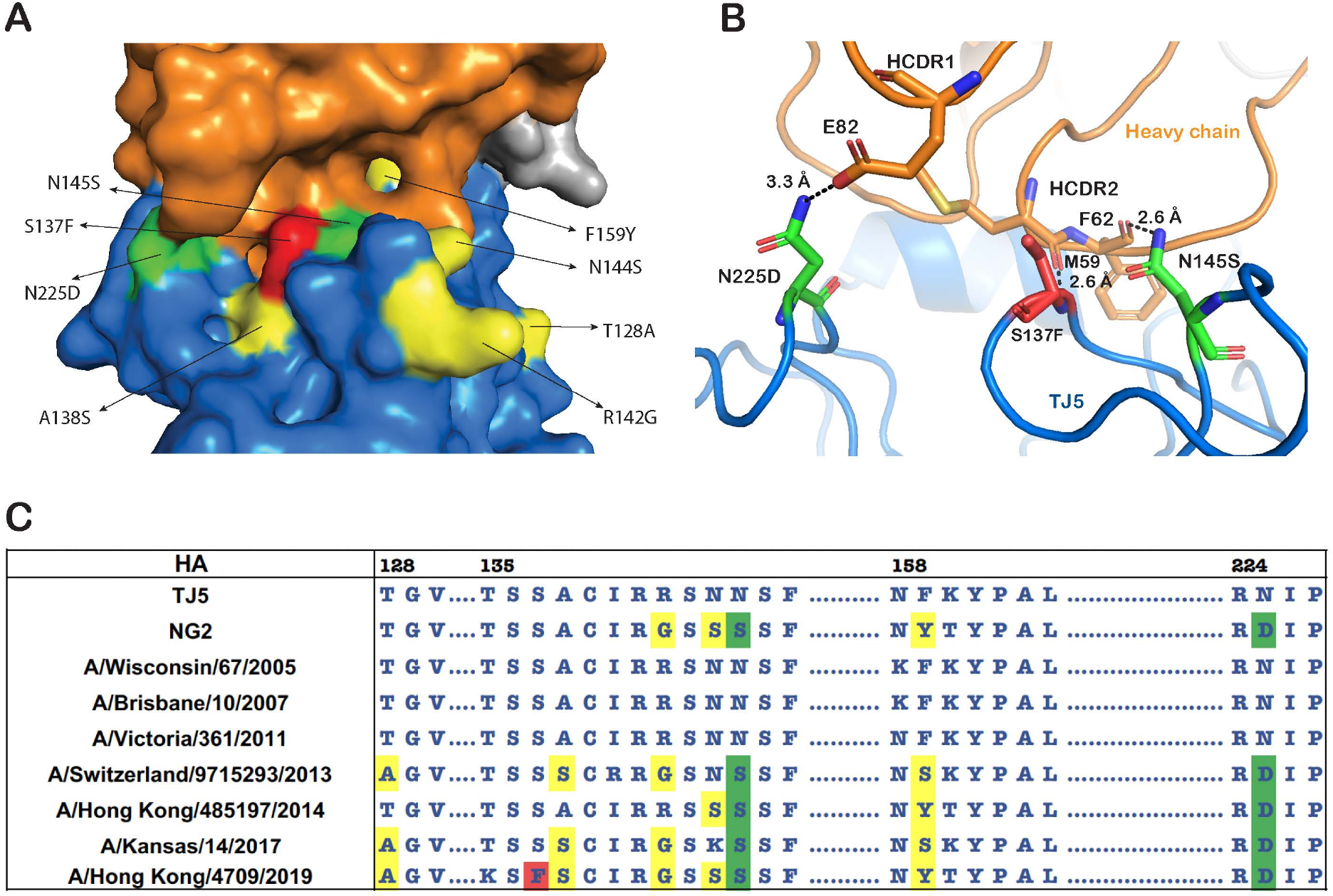
The amino acid mutations in HA binding site decrease mAb binding. A) HA mutations found near binding site in NG2, SWZ13 and HK19 (yellow). Green shows the mutations that are essential to recognition by TJ5-5. Red shows a mutation specific to HK19 essential to recognition. B) Shows the interactions between the HC and HA formed by the residues affected by mutations in NG2, SWZ13 and HK19 N145S, N225D (green) and S137F (Red). The heavy chains are shown as orange, the light chains are shown in grey and the HA is shown in blue. C) Alignment of HA sequences from H3 COBRAs TJ5 and NG2 and seasonal H3N2 strains. The mutations located near the RBS are highlighted in yellow and mutations essential for binding are highlighted in green (N145S, N225D) and red (S137F).

## Discussion

HA is the most abundant glycoprotein on the influenza virus surface, promoting viral entry by engaging the receptor binding site on the head domain via sialic acid receptors on host cells, and mediating membrane fusion between the virus and host cell membranes through the fusion peptide on the stalk domain. Antibodies targeting the HA protein that block hemagglutinin activity or prevent HA fusion activity are protective against infection, but this protection is often strain or clade-specific. The COBRA methodology incorporates HA sequences of past and current circulating seasonal influenza strains to develop a vaccine that elicits more broadly neutralizing antibodies. As nearly all individuals have pre-existing immunity to influenza viruses, COBRA HA vaccines will likely recall memory B cells from previous influenza infection and vaccination, in addition to generating de novo B cell responses. To determine the antigenic features of H3 COBRA HA proteins and to assess potential recall patterns, we probed B cell recognition of three H3 COBRA HA proteins using samples from subjects vaccinated with the split-inactivated 2017-2018 seasonal influenza vaccine. TJ5 and J4 COBRA HA proteins were recognized at higher frequencies than the NG2 COBRA HA, likely due to NG2 COBRA HA incorporating sequences from later vaccine years than TJ5 and J4 proteins.

To further assess this differential recognition, we generated twelve H3 COBRA reactive monoclonal antibodies (mAbs) from the human subjects, and we characterized mAb binding avidity and functional activity using hemagglutination inhibition (HAI) and focal reduction assays. We also determined target mAb epitopes on COBRA antigens. In addition to their broad binding properties, the isolated mAbs displayed functional activity in HAI and neutralization assays. mAbs TJ5-4, TJ5-5, TJ5-6, TJ5-8, TJ5-9, TJ5-13, and TJ5-15 had the broadest and most potent neutralizing activity against a panel of seasonal H3N2 viruses. Competitive biolayer interferometry experiments showed that all mAbs target the RBS on the head domain except for mAb TJ5-1, which was found to target the stalk domain, and as expected, this mAb has the broadest neutralizing activity.

To determine the differential recognition of TJ5/J4 COBRA HAs versus the NG2 COBRA HA by pre-existing human B cells, we utilized a representative mAb, TJ5-5, which had matching binding data to our oligoclonal B cell analysis, specifically, poor recognition of COBRA NG2 and high avidity for TJ5 and J4. A cryo-EM structure of mAb TJ5-5 in complex with COBRA TJ5 identified the presence of amino acid substitutions (N145S and N225D) at the mAb binding site of TJ5 that would likely result in the diminished recognition of NG2. As further evidence of the importance of these substitutions, mAb TJ5-5 had weaker neutralizing activity against A/Kansas/14/2017 and A/Switzerland/9715293/2013 viruses bearing these substitutions, but retained potent neutralizing activity against A/Wisconsin/67/2005, A/Brisbane/10/2007 and A/Victoria/361/2011 viruses that do not have these substitutions. Additionally, our structure verified the structural integrity of the TJ5 HA protein, which is the first structure of an H3 COBRA HA protein reported.

In conclusion, this study provides evidence that H3 COBRA HA proteins can induce the recall of B cells elicited by seasonal influenza vaccination, and verifies the antigenic and structural features of the TJ5 COBRA HA. One drawback of our study is that we utilized B cells for our analysis collected 21-28 days following seasonal influenza vaccination, and that the B cell responses and mAbs may partially reflect responses from previous influenza seasons rather than just the 2017-2018 season. Importantly, our data suggest that a small number of amino acid substitutions in next-generation COBRA HA proteins can have a profound effect on pre-existing B cell recognition, and while COBRA HA proteins indeed elicit broader antibody responses, consistent monitoring of these responses must be assessed as influenza viruses evolve to ensure protection is maintained.

## Acknowledgements

This work was supported by the Collaborative Influenza Vaccine Innovation Centers (CIVIC) contract by the National Institute of Allergy and Infectious Diseases, a component of the NIH, Department of Health and Human Services, under contracts 75N93019C00052 (T.M.R., J.J.M.). J.J.M. is partially supported by National Institutes of Health grant K01OD026569. T.M.R is supported as an Eminent Scholar by the Georgia Research Alliance. Cryo-EM data was collected at the Hauptman-Woodward Institute Cryo-Electron Microscopy Center.

## Data availability

The 3D reconstruction of the TJ5 + TJ5-5 structure was deposited to the PDB under accession ID 7TZ5 and to EMDB under accession ID EMD-26202.

---

**Table S1.**
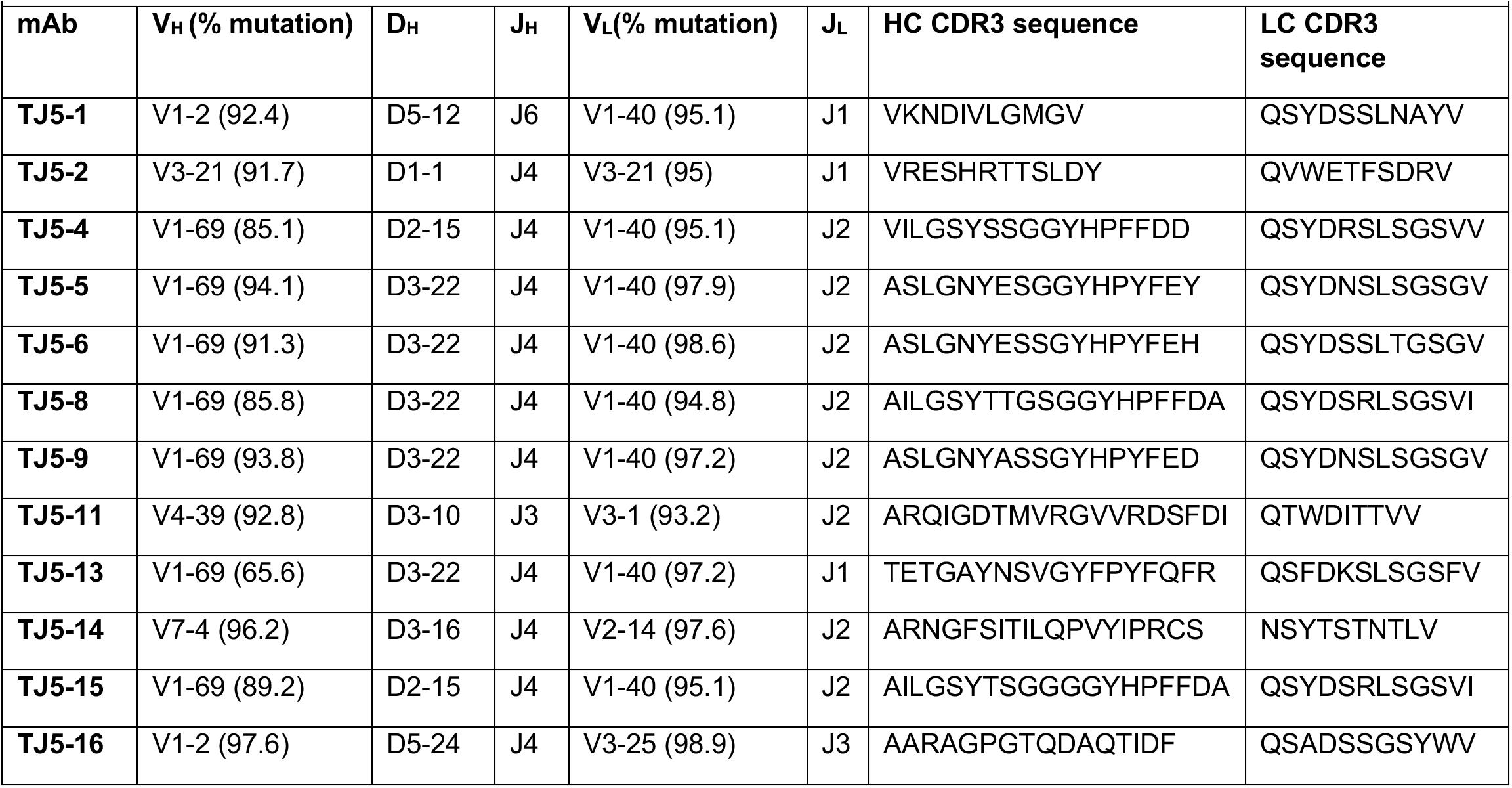
Monoclonal antibody sequence determinants.

**Table S2.**
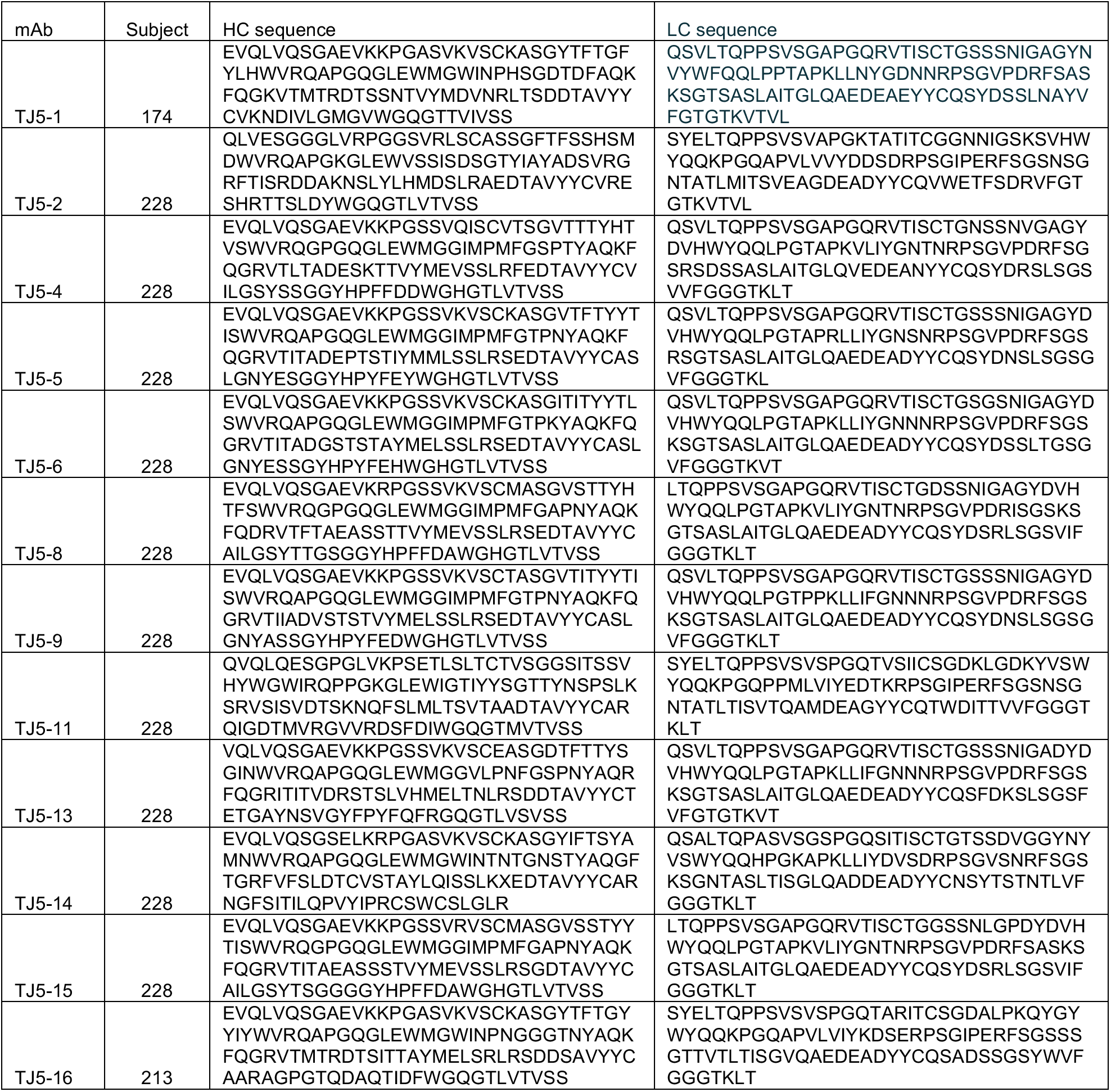
Monoclonal antibody sequences.

**Figure S1.**
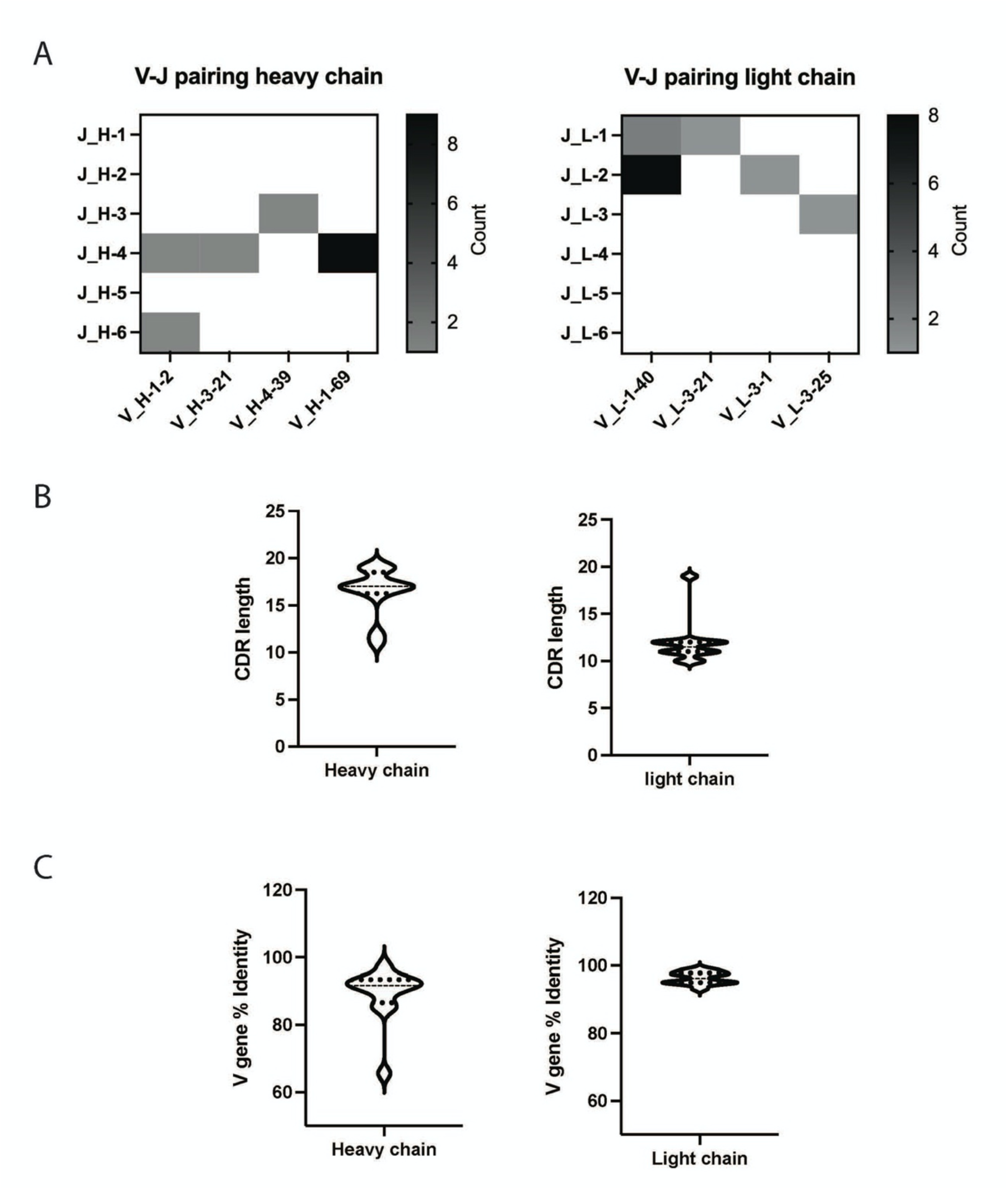
V-J analysis of H3 COBRA reactive mAbs. A) V-J pairings utilized by heavy and light chains. V genes used are on the X-axis and J genes are on the Y-axis. The grey scale counts the number of mAbs using each V-J pairing. B) The range of lengths of heavy and light chain junctions in each isolated mAb. C) The percent identities of the variable genes to the germline sequence in the heavy and light chains of isolated mAbs.

**Table S3.** EM data collection information and statistics.

| **Table S3. EM data collection information and statistics** |  |
| --- | --- |
| **EM data collection** |  |
| Sample | TJ5 HA + TJ5-5 |
| PDB ID | 7TZ5 |
| EMDB ID | 26202 |
| Microscope | Glacios |
| Voltage (kV) | 200 |
| Detector | Falcon 4 |
| Magnification (nominal) | 190,000 |
| Pixel size (Å/pix) | 0.526 |
| Exposure rate (e<sup>-</sup>/Å<sup>2</sup>) | 57.22 |
| Number of frames | 30 |
| Defocus range (µm) | 0.9-2 |
| Micrographs collected | 10872 |
| Micrographs used | 10872 |
| Particles extracted (total) | 887,924 |
| **3D reconstruction statistics** |  |
| Particles | 101267 |
| Symmetry | C3 |
| Map sharpening B-factor | -146.5 |
| Unmasked resolution at 0.5 FSC (Å) | 3.9 Å |
| Masked resolution at 0.5 FSC (Å) | 3.6 Å |
| Unmasked resolution at 0.143 FSC (Å) | 3.4 Å |
| Masked resolution at 0.143 FSC (Å) | 3.2 Å |
| **Model refinement and validation statistics** |  |
| Refinement package | Phenix |
| Refinement tool | Real-space refinement |
| Refinement strategies | Min global, local_grid search, adp, ss restraints, Ramachandran restraints |
| Atoms | 16023 |
| Residue | Protein: 2142 |
| Bond length (Å) | 0.009 |
| Bond angles (°) | 1.396 |
| MolProbity score | 2.26 |
| Clash score | 15.18 |
| Ramachandran plot (%) Outliers | 0.42 |
| Ramachandran plot (%) Allowed | 10.62 |
| Ramachandran plot (%) Favored | 88.95 |
| Rotamer outliers (%) | 0.36 |
| Cβ outliers (%) | 0.46 |
| CaBLAM outliers (%) | 5.70 |
| ADP (B-factors) protein | 89.34 |
| CC (mask) | 0.83 |

**Figure S2.**
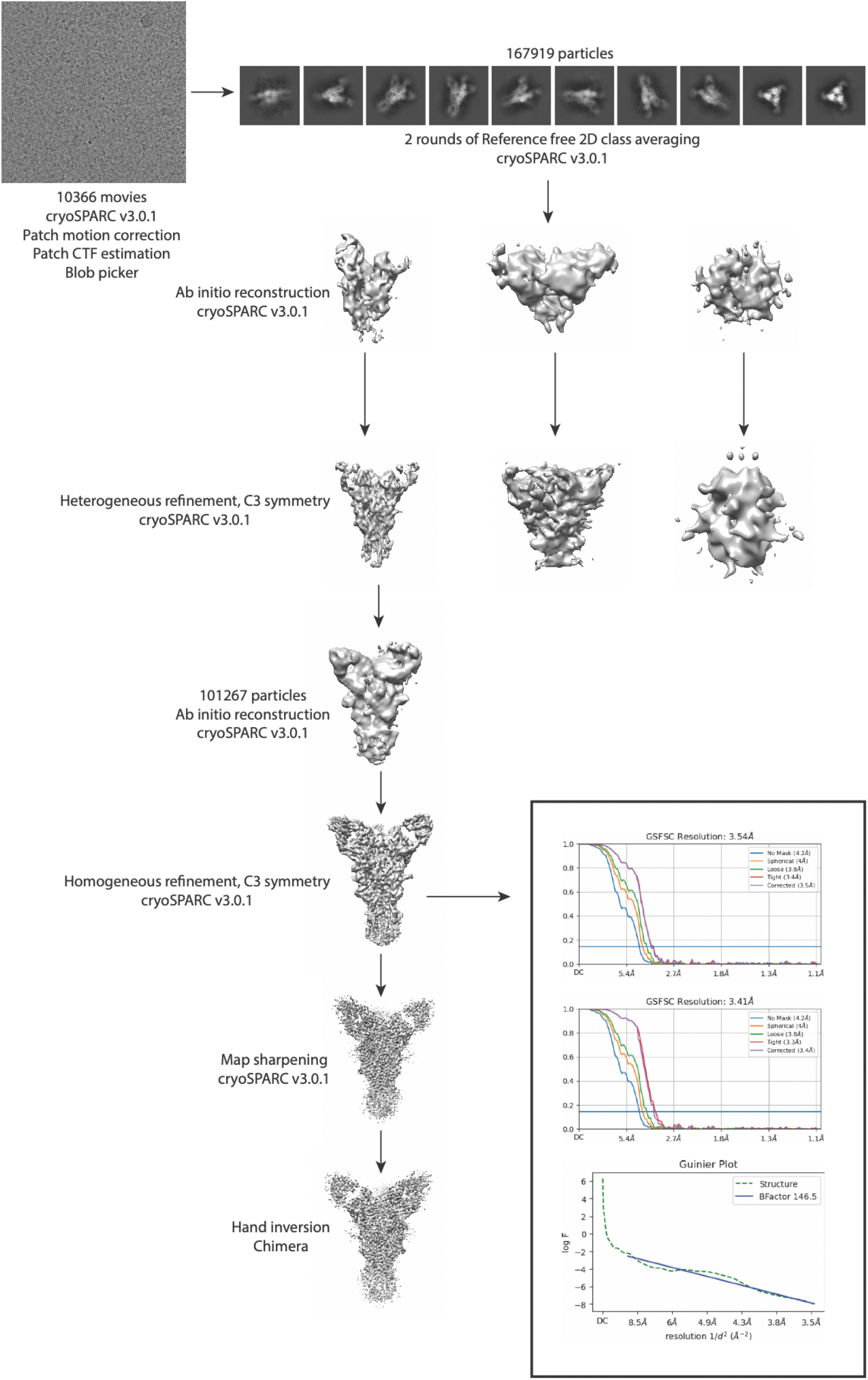
Cryo-EM processing workflow.

